# IκBζ is constitutively expressed in human and murine airway epithelium

**DOI:** 10.1101/255331

**Authors:** Kruthika Sundaram, Srabani Mitra, Sebastian Lorscheid, Haley Steiner, Anasuya Sarkar, Konstantin Shilo, Prosper N. Boyaka, Klaus Schulze-Osthoff, Mark D. Wewers

## Abstract

**AIM:** IκBζ is a transcriptional factor induced in immune cells upon Toll-like receptor (TLR) activation. Recent studies demonstrate unconventional, constitutive expression of IκBζ in epithelium of mouse skin and eyes, possibly reflecting continuous activation of TLRs by pathogen-associated molecular patterns (PAMPs). In this context, the lung epithelium which constitutes another important barrier also expresses IκBζ but may not be as actively exposed to pathogens as skin and eyes. Our aim was to determine if IκBζ expression in the lungs is constitutive or induced.

**SIGNIFICANCE:** IκBζ is linked to lung disorders due to its role in regulating protective cytokines and antimicrobial peptides in airway epithelium and can therefore be a potential biomarker and a key therapeutic target.

**METHODS:** We evaluated IκBζ expression in airway epithelia of healthy humans and three kinds of mice: normal, gnotobiotic and *Nfkbiz^−/−^* knockout, using immunostaining and immunoblotting.

**RESULTS:** Immunohistochemistry of ciliated airway epithelial cells in healthy humans and normal mice was positive for IκBζ. The pathogen free airway cells from gnotobiotic mice also stained positive, suggesting that lung epithelial IκBζ expression does not require induction by PAMPs. Although lung epithelia from *Nfkbiz^−/−^* knockout mice also stained positive, this knockout may not have eliminated exons 3-4 and 9-14, and so did not provide the specificity control for the IκBζ antiserum. Importantly, immunoblotting tissue homogenates from gnotobiotic mouse lungs and primary human airway epithelial cells demonstrated constitutive IκBζ expression at its correct 86 kDa size.

**CONCLUSIONS:** Our data demonstrates constitutive expression of IκBζ protein in airway epithelium, indicating a potential role for this molecule in lung homeostasis.

## BACKGROUND

IκBζ, also known as MAIL (molecule with ankyrin repeats induced by LPS) or INAP (IL-1-inducible nuclear ankyrin-repeat protein), is a protein encoded by the gene *NFKBIZ* (1–3). It predominantly functions as a transcription factor to regulate the expression of many downstream molecules that are involved in host defense, such as IL-6, IL-12 (p40), granulocyte-macrophage colony-stimulating factor, neutrophil gelatinase-associated lipocalin, human β defensin 2 and monocyte chemoattractant protein 1 (4–9). IκBζ is a homolog of the IκB group of proteins, due to the presence of an ankyrin repeat region in its carboxy-terminus, with the help of which it binds to NFκB p50 homodimers, typically inside the nucleus of the cell (1, 3). The amino-terminus of IκBζ encodes a transcriptional activation domain and a nuclear localization sequence. There are two known isoforms of the protein: long (IκBζ-L) and short (IκBζ-S), of which IκBζ-L is predominantly expressed at the protein level (1). Apart from its role as a transcription factor, IκBζ is known to exhibit other functions. For instance, IκBζ modulates chromatin remodeling, by selectively regulating H3K4 trimethylation and the assembly of the pre-initiation complex at the promoter of downstream genes (10).

A majority of studies have described IκBζ as a primary, early response gene induced rapidly following TLR stimulation of immune cells including mononuclear phagocytes, NK cells, T and B lymphocytes (6, 8, 11). Nevertheless, constitutive expression of the *NFKBIZ* gene has been demonstrated in mucosal and epidermal sites that interact with the outside environment (12, 13), indicating the possibility of chronic exogenous challenges inducing stable IκBζ expression in these cells. In support of this concept, IκBζ knockout mice display severe inflammation of the epithelium lining their eyes and skin (13–15), suggesting the importance of IκBζ in maintaining immune homeostasis of the host. Interestingly, IκBζ has been demonstrated to regulate the expression of pro-inflammatory cytokines and anti-microbial peptides in airway epithelial cells that line the respiratory mucosal surface, in mammalian models of asthma and pneumonia (4, 5, 16). Studies have shown baseline *NFKBIZ* gene expression in lung tissues (3, 14), although the induction of IκBζ protein expression in the airway epithelium in response to environmental agonists remains understudied. The present study therefore explored the expression of IκBζ in both human and mouse airway epithelial cells. Using immunohistochemistry (IHC), we detected positive staining in the nuclei of ciliated epithelial cells in the upper airway tract of human and wildtype gnotobiotic mice. Immunoblotting of tissue homogenates from gnotobiotic mice and primary human airway epithelial cells demonstrated constitutive expression of IκBζ. Thus, our results support a role for IκBζ in airway protection.

## MATERIALS AND METHODS

### Reagents and antibodies

The reagents were obtained from the following sources: IL-1 receptor antagonist, IL-1Ra (Amgen), penicillin-streptomycin (Invitrogen), fetal bovine serum FBS (Atlanta Biologicals), Dulbecco modified Eagle medium low glucose DMEM (Invitrogen), Bronchial Epithelial Growth Medium BEGM bullet kit (Lonza), Bronchial Air Liquid Interface B-ALI media (Lonza), Small airway epithelial cell media and detach kit (PromoCell), recombinant human IL-1β was purchased from R&D systems. Rabbit antiserum against IκBζ was generated in our laboratory using recombinant protein expressed in *Escherichia coli* (8) (17). Commercial antibodies against IκBζ were purchased from Thermo Fisher Scientific (PA5-52703), Sigma Aldrich (HPA010547), Abcam (ab221914) and LS Biosciences (LS-C294627). Scrambled siRNA control and siIκBζ (sequence UGAUGGACCUGCUUGCAAA) were purchased from Dharmacon Thermo Scientific. Lamin B1 antibody and beta-actin antibody (monoclonal clone C4) were purchased from Santa Cruz Biotechnology and MP Biomedicals respectively.

### Mouse lung tissue

Lungs were harvested from wildtype B10RIII mice weighing 17-20 grams and lungs and tracheae were isolated from 9-12 weeks old gnotobiotic male Swiss Webster mice, weighing 17-21 grams. Wildtype and *Nfkbiz^−/−^* knockout mice were allowed free access to food and water, while the gnotobiotics were bred and maintained in sterile isolators. All animal experiments were performed according to animal protocols approved by the Animal Care Use Committee of the Ohio State University College of Medicine. Lungs from wildtype mice were equivalently inflated with an intratracheal injection of a similar volume of 4% paraformaldehyde solution (Sigma) to preserve the pulmonary architecture. The lungs were then embedded in paraffin. The sections of all organs collected were then stained with hematoxylin and eosin or with anti-IκBζ antiserum, along with appropriate controls.

### Human lung tissue

Anonymized human lung tissue slides were provided by the Ohio State University Department of Pathology, Tissue Archive Service. Other investigators may have received specimens from the same subjects.

### Immunohistochemistry

The lung tissue slides were labeled with IκBζ antiserum at the specified dilution with anti-rabbit secondary antibody at a 1:1000 dilution at the Comparative Pathology and Mouse Phenotyping Shared Resource, at the Veterinary school of OSU.

### Cell culture

BEAS2B and HeLa cells (purchased from ATCC) were maintained in DMEM low glucose, supplemented with 10% FBS and 1% penicillin-streptomycin, in a 37°C humidified incubator with 5% CO_2_. Monocytes were obtained from fresh buffy coats purchased from the American Red Cross. Monocytes were purified from blood by Histopaque density gradient centrifugation using lymphocyte separation media (Cellgro) followed by CD14 positive selection as described previously (8). Purified monocytes and monocytic cell line THP-1 were both maintained in RPMI1640 supplemented with 10% FBS. Primary human bronchial epithelial cells (HBECs), purchased from Lonza were allowed to differentiate in B-ALI media in 24-well inserts (6.5 mm) as per manufacturer’s instructions. HBECs (5×10^4^ cells/33mm^2^) were stimulated with rhIL-1β (10ng/ml) for 3 hours, with or without 30 minutes of pretreatment with IL-1Ra (10ug/ml). Cells are then lysed and immunoblotted.

### Generation of recombinant IκBζ

IκBζ-S was cloned in pET32a bacterial vector with an N-terminal thioredoxin tag and a C-terminal histidine tag and then over expressed in BL21DE3RIL strain of bacteria. The over expressed protein was purified using HisPur cobalt resin and dialyzed into a 20mM Tris buffer, pH 7.5 with 150mM NaCl and 1mM beta mercaptoethanol, using a Slide-a-lyzer, both from Thermo Scientific. Expression of purified rIκBζ-S was confirmed using immunoblotting with both anti-IκBζ antiserum and His antibody.

### Blocking non-specißcity of anti-IκBζ antiserum using THP-1 extracts

Unstimulated THP-1 cells (~7×10^7^ cells) that do not express IκBζ were washed to remove culture media and re-suspended in 10ml of homogenization buffer (0.25M sucrose, 1mM EDTA, 20mM HEPES pH 7.5) with protease inhibitors mixture (Sigma-Aldrich). The cells were then dounced on ice and centrifuged at low speed (~100rpm) to remove unbroken cells. The supernatant was centrifuged at high speed (~2000rpm) and was separated into the cellular extract (CE) fraction and the intact nuclei. The intact nuclei that pellet out were then re-suspended in 1ml of Buffer W (20mM HEPES pH 7.5, 10mM KCl, 1.5mM MgCl2, 1mM EGTA, 1mM EDTA) with protease inhibitors, sonicated on ice for 10-15 minutes and centrifuged at high speed to yield the nuclear pellet (Nu) and the nuclear extract (NE) that was then used for blocking. Our laboratory made anti-IκBζ antiserum was incubated with 1000-2000 times excess by volume of the THP-1 NE at room temperature for 1-2 hours. This antigen-antibody mixture was then used for immunoblotting or immunostaining.

### Immunostaining of BEAS2B cells

BEAS2B cells (10^5^ cells/ml) were seeded onto sterile coverslips placed inside 12 well plates and cultured overnight. The cells were then stimulated with IL-1β (10ng/ml) for 3-4 hours.

The media was removed and ice cold 100% methanol was slowly added to the coverslips to fix the cells. The cells were left in methanol at 4°C overnight. The cells were then immunostained using the Vectastain Elite ABC kit from Vector laboratories, as per manufacturer’s instructions. The primary antibody used was either anti-IκBζ antiserum alone at 1:2000 dilution or with anti-IκBζ antiserum incubated with excess THP-1 NE.

### Immunoprecipitation

Human monocytes were plated in 10 ml petri dishes and were stimulated with 1 μg/ml LPS for 3 h. The cells were then lysed in RIPA buffer (50mM Tris [pH 7.5], 1% sodium deoxycholate, 1mM EDTA, 1mM NaF, 150mM NaCl, 1% NP-40) with the protease inhibitor cocktail mix, 10 mM methoxysuccinyl-Ala-Ala-Pro-Val-chloromethyl ketone, 1 mM phenylmethylsulfonyl fluoride (Sigma-Aldrich), and 3 mM sodium orthovanadate. Cell lysates were pre-cleared using protein A beads (Invitrogen). Pre-cleared lysates were immunoprecipitated using anti-lamin B1 antibody and these immunoprecipitated samples were incubated with protein A-sepharose beads (Invitrogen) for 2h at 4°C, followed by centrifugation at 16,000 × *g* for 5 min and washing. The proteins were eluted with 2X SDS-PAGE sample buffer at 95°C for 10 min and were separated by SDS-PAGE and blotted with anti-lamin B1 antibody and anti-IκBζ antiserum.

### 2D gel analysis

IL-1β stimulated HeLa cells were immunoprecipitated with anti-IκBζ antiserum as described above. The immunoprecipitated samples were run on a 2D gel electrophoresis system at the OSU’s Campus Chemical Instrument Center (CCIC) mass spectrometry and proteomics core facility, on an 18-cm 4-7 IPG strip, since the predicted isoelectric point for IκBζ was around 6. The separated proteins were then either transferred onto a PVDF membrane and immunoblotted with anti-IκBζ antiserum or were further processed for mass spectrometry analysis.

### Mass spectrometry

The 2D gel was stained with Coomasie blue and bands corresponding to 86 kDa (the size of IκBζ-L) and 68 kDa (the size of IκBζ-S) were excised and submitted to OSU’s proteomics facility. Samples underwent automated in-gel protein digestion in the MassPrep station. Briefly, they were destained, reduced and incubated with iodoacetamide in ammonium bicarbonate for 20 min at 37 °C. They were then washed with ammonium bicarbonate/water and dehydrated with acetonitrile. The extracted proteins were in-gel digested with 6 ng/μl trypsin in 50 mM ammonium bicarbonate for 5 h at 37 °C. Mass spectra of resulting peptides were recorded on the ESI-TRAP spectrometer. The resulting peptides were matched with their corresponding proteins with ProFound by searching the non-redundant data base maintained at the NCBI (www.ncbi.nlm.nih.gov). The cut off for ion score was kept at 20, where the score was calculated as −10*Log(P), and P, the probability that the observed match is a random event. The following parameters were used for the search at Pro-Found: taxa *Homo sapiens*, fixed modifications carbamidomethyl, variable modifications oxidation, allowed incomplete cleavages 2, monoisotopic masses, and a peptide mass tolerance of ±1.8 Da.

### Tissue homogenization

Tissue chunks of lungs or trachea from mice were added to SDS lysis buffer (100 mM NaCl 500 mM Tris, pH 8.0 10% SDS) and vortexed for about 30 minutes till the chunks dissolved. The tissues were then syringed approximately 20 times and boiled at 100°C for complete extraction. The tissue extracts were then protein estimated and immunoblotted for IκBζ expression.

### Small interfering RNA (siRNA) mediated knockdown of IκBζ

BEAS2B cells were transfected with 100pmol of scrambled siRNA control or siRNA specific to IκBζ using lipofectamine (Invitrogen). After 20 hours, media was replaced, following which the cells were stimulated with rIL-1β (10ng/ml) for 4 hours.

### Preparation of cell lysates and immunoblotting

Cells were lysed in TN1 lysis buffer (50 mM Tris-HCl, pH 8.0, 125 mM NaCl, 10 mM ethylene diaminetetraacetic acid (EDTA), 10 mM NaF, 10mM sodium pyrophosphate, 1% Triton X-100) with the protease and phosphatase inhibitor cocktails. The extracts were incubated on ice for 15 minutes, syringed 10-15 times and centrifuged at 16,000g for 10 minutes. The lysates were transferred to a new Eppendorf tube and total protein in each sample was determined using Lowry assay (Bio-Rad). Equal protein (20ug per lane) was loaded onto a NuPAGE 7% Tris-acetate gel (Invitrogen). The separated proteins were transferred onto polyvinylidene difluoride membranes. The membranes were blocked with 10% nonfat milk, incubated overnight at 4°C with the primary antibody, washed and then stained with appropriate peroxidase-conjugated secondary antibody for an hour. The protein bands were visualized by enhanced chemiluminescence substrate (GE Healthcare) by autoradiography.

## RESULTS

### Anti-IκBζ antiserum detects IκBζ

The first step to address the aim of the study was to find an antibody that would detect IκBζ with high sensitivity and specificity. We started by validating the anti-IκBζ antiserum generated in our laboratory. To do this, we transfected BEAS2B cells with siRNA specific to IκBζ and stimulated them with rIL-1β for 4 hours. The anti-IκBζ antiserum detected the expected pattern of IκBζ-L expression at the correct size of 86 kDa. The protein was induced in response to IL-1β stimulation which was blocked in the presence of si-IκBζ (**Figure S1A**). However, the antiserum also detected non-specific proteins. We then tested a panel of commercial antibodies from different manufacturers. None of them detected IκBζ with the same sensitivity as our laboratory generated anti-IκBζ antiserum (**Figure S1B**). Additionally, they also detected nonspecific bands. We hence used our anti-IκBζ antiserum for all further experiments in this study.

### IκBζ staining observed in mammalian lung tissue

To understand IκBζ’s behavior in airway epithelial cells, we immunostained lung tissue slices from normal human subjects (**Figure 1A**) and from wild type mice (**Figure 1B**) with the anti-IκBζ antiserum. We observed a significant increase in staining with anti-IκBζ antiserum compared to the preimmune control serum and the secondary antibody negative control. This signal was predominantly present in the nuclei of ciliated epithelial cells lining the central airways, which was the expected cellular location for IκBζ due to its known role in transcription regulation.

**Figure 1.**
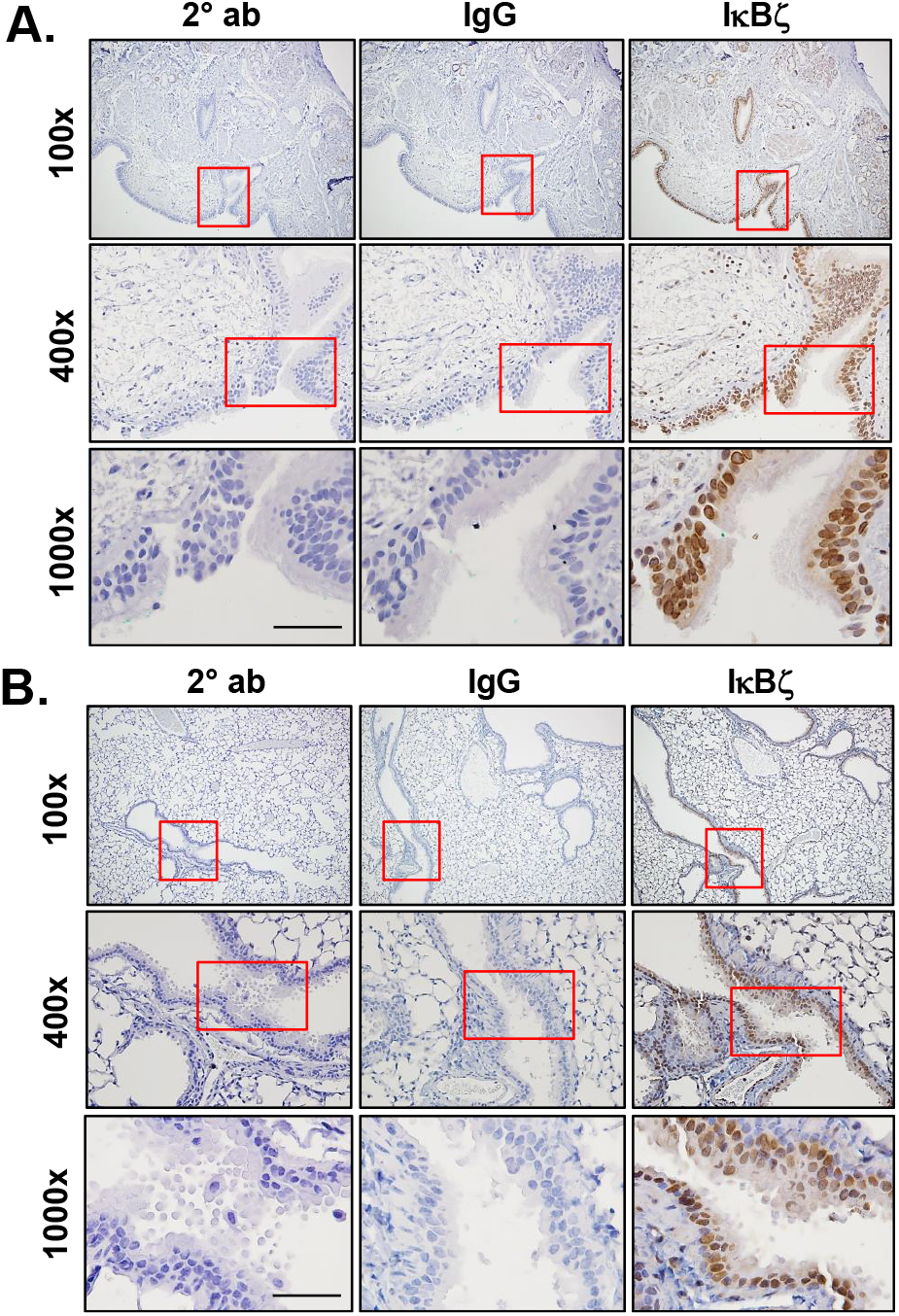
IκBζ expression in normal lung tissue. Lung tissue slices from **(A)** healthy human individuals were immunostained with secondary antibody (2° ab) alone (1:1000), rabbit control IgG (1:1000), or anti-IκBζ antiserum (1:1000) **(B)** wild type normal mice were immunostained with secondary antibody alone (1:1000), IgG isotype control (1:2000), or anti-IκBζ antiserum (1:2000), observed at 100, 400 and 1000X magnifications. The scale bar denotes 50μm for the 1000X magnification panels. The results are representative of 3 different mice/humans. Red boxes represent regions that were captured at higher magnifications.

### Gnotobiotic lungs stain positive for IκBζ expression

Because IκBζ is considered to be a classic example of an early response gene, we asked if the baseline IκBζ expression was due to exposure to bacterial PAMPs. Therefore, we examined lung tissue from gnotobiotic (germ-free) mice. Surprisingly, the ciliated epithelial cells of the gnotobiotics also immunostained for IκBζ expression, both in the major bronchi (**Figure 2A**) and trachea (**Figure S2**), suggesting that expression of IκBζ in the airway epithelium does not require induction by bacterial PAMPs.

**Figure 2.**
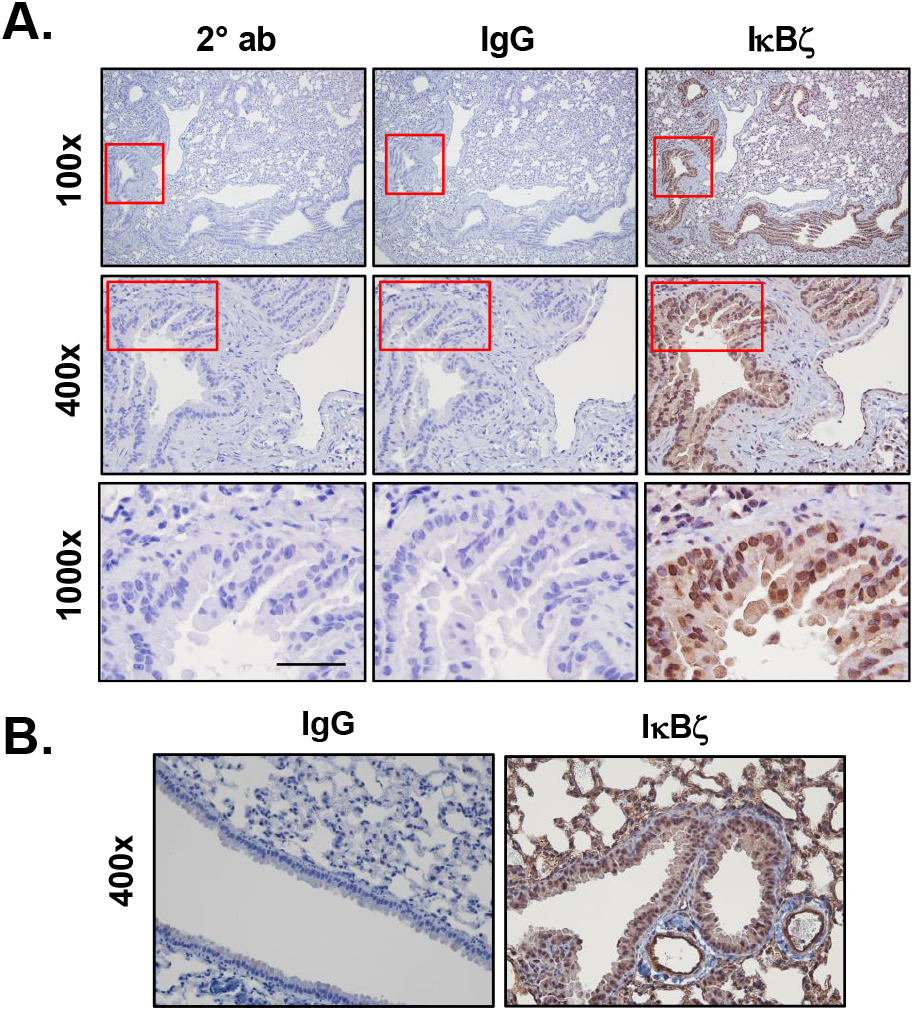
IκBζ expression in lung tissue from gnotobiotic and *Nfkbiz^−/−^* knockout mice. Lung tissue slices from **(A)** gnotobiotic mice were immunostained with secondary antibody (2° ab) alone (1:1000), IgG isotype control (1:1500), or anti-IκBζ antiserum (1:1500), observed at 100, 400 and 1000X magnifications and from **(B)** *Nfkbiz^−/−^* knockout mice were immunostained with IgG isotype control (1:2500), or anti-IκBζ antiserum (1:2500), observed at 400X magnification. The scale bar denotes 50μm for the 1000X magnification panels. The results are representative of 3 different mice. Red boxes represent regions that were captured at higher magnifications.

### Anti-IκBζ antiserum also detects lamin B1

To further validate these results, we immunostained lung tissue samples from *Nfkbiz^−/−^* knockout mice (**Figure 2B**). Unexpectedly, we obtained a positive signal. However, this knockout (13) may not have eliminated exons 3-4 and 9-14, and so did not provide the specificity control for the IκBζ antiserum.

In an attempt to prove specificity of the antiserum, we used column-purified recombinant IκBζ-S expressed in bacterial cells (**Figure 3A**) to block IκBζ detection by anti-IκBζ antiserum. Purified rIκBζ-S, when incubated in excess with anti-IκBζ antiserum, blocked not only the IκBζ-L 86kDa band but also all of the nonspecific bands (**Figure 3B**), as observed previously in our group (17). This result therefore still required further specificity confirmation.

**Figure 3.**
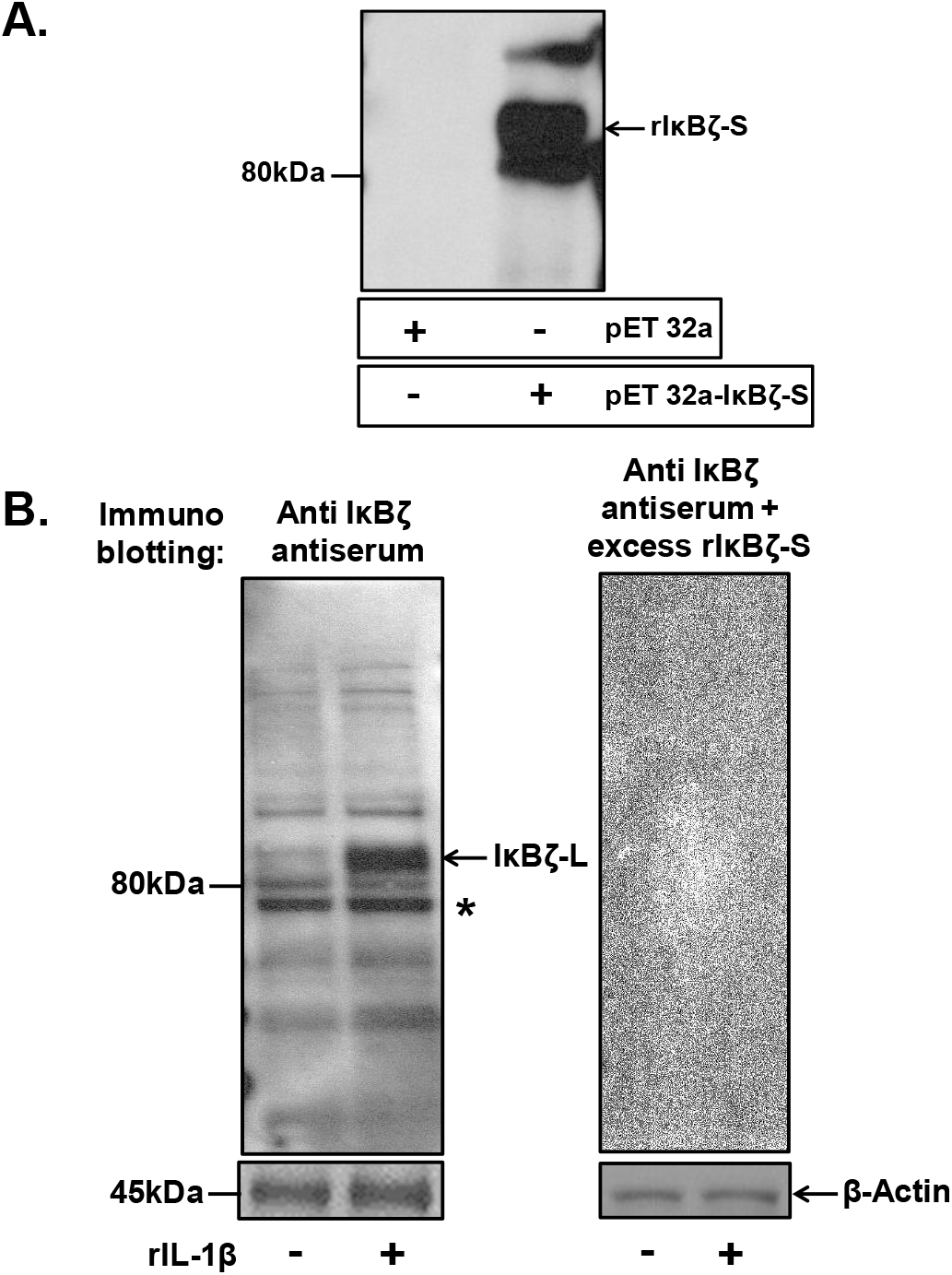
rIκBζ blocks non-specific signal detected by anti-IκBζ antiserum. **(A)** Immunoblot of column purified rIκBζ-S detected by anti-IκBζ antiserum (1:2000). It runs at ~82 kDa due to the presence of His and thioredoxin tags. **(B)** IL-1β-stimulated BEAS2B cell extracts were immunoblotted with anti-IκBζ antiserum alone or with anti-IκBζ antiserum incubated with excess rIκBζ-S protein. Asterisk represents non-specific protein detected. Beta actin was used as protein loading control. The results are representative of 3 independent experiments.

To evaluate what other proteins were detected by the anti-IκBζ antiserum, we performed 2D electrophoresis on samples obtained by immunoprecipitation using the anti-IκBζ antiserum from lysates of HeLa cells stimulated with IL-1β (**Figure 4A**). Mass spectrometry analysis of the cluster of spots identified IκBζ-L at the expected size of 86 kDa towards the acidic end of the pH strip. Interestingly, spots that ran at 68 kDa with isoelectric points close to neutral pH were identified as the nuclear protein lamin B1 and not IκBζ-S which also runs at 68 kDa. The cross reactivity with lamin B1 was further confirmed when lamin B1 was detected by anti-IκBζ antiserum in the sample immunoprecipitated with anti-lamin B1 antibody from lysates of monocytes stimulated with LPS (**Figure 4B**).

**Figure 4.**
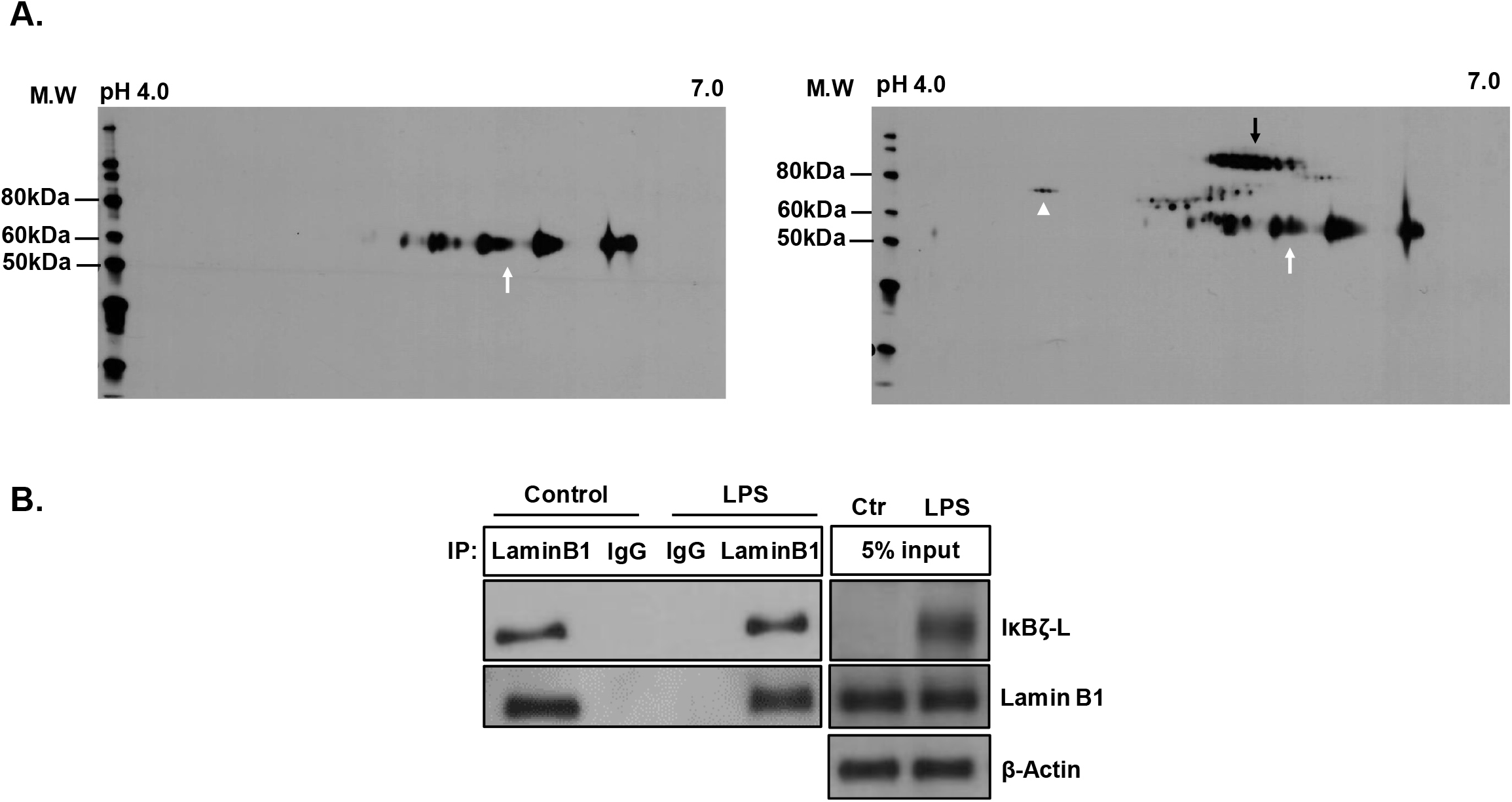
Anti-IκBζ antiserum detects lamin B1 protein. **(A)** Lysates of HeLa cells stimulated with IL-1β were immunoprecipitated using anti-IκBζ antiserum (right panel) or IgG control (left panel) and run on 2D SDS-PAGE, followed by immunoblotting with anti-IκBζ antiserum (1:2000). White arrow head represents lamin B1 while black arrow represents IκBζ-L, as identified by ESI-TRAP mass spectrometry. White arrow represents IgG heavy chain. **(B)** Lysates of monocytes stimulated with LPS were immunoprecipitated using anti-lamin B1 antibody and blotted with respective antibodies as indicated. The results are representative of 3 independent experiments.

Since our attempts to specifically block IκBζ detection alone by preblocking our anti-IκBζ antiserum with recombinant IκBζ failed, we tried to remove all the non-specific IgG molecules in our anti-IκBζ antiserum, leaving the IgGs specific to IκBζ available for binding. To do this, we incubated anti-IκBζ antiserum with an excess of nuclear extracts (NE) from unstimulated THP-1 cells that contained lamin B1 but no IκBζ, i.e., the non-specific antigens detected by the anti-IκBζ antiserum (**Figure S3A**) (8). Using this “precleared” anti-IκBζ antiserum on IL-1β activated BEAS2B cells significantly blocked the immunoblot and immunostaining for lamin B1 (**Figure S3B & C**) without reducing the IκBζ signal. However, THP-1 extract did not completely block the non-specific IgGs in the anti-IκBζ antiserum, despite its presence in huge excess, as seen in the western blot and the immunostain (**Figure S3B & top panel of S3C**).

### Immunoblotting demonstrates definitive constitutive IκBζ expression in lung epithelium

We then resorted to immunoblotting with anti-IκBζ antiserum to perform size-based detection of specific IκBζ expression and thus definitely demonstrate its constitutive expression in lung epithelium. Tissue homogenates from gnotobiotic mice showed IκBζ-L expression at the correct molecular size (~86 kDa) both in the lungs and the trachea (**Figure 5A**). We then compared the expression of the protein in differentiated primary human bronchial epithelial cells (HBEC). HBECs displayed baseline IκBζ expression in the untreated controls (**Figure 5B**) that was further induced by IL-1β and this induction was inhibited in the presence of IL-1 receptor antagonist (IL-1Ra).

**Figure 5.**
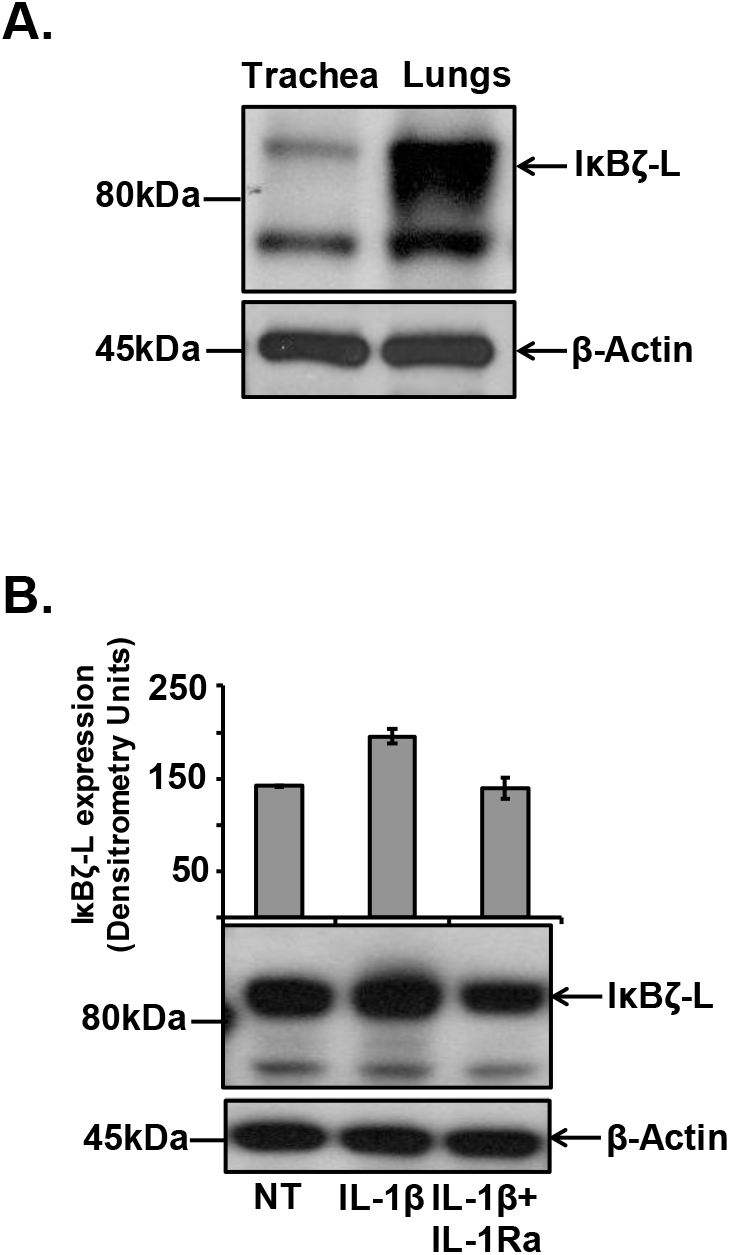
Immunoblotting demonstrates constitutive IκBζ expression in airway epithelium. **(A)** Immunoblot of homogenates from trachea and lungs of gnotobiotic mice, detected by anti-IκBζ antiserum. The results are representative of 3 different mice. **(B)** HBECs (2×10^5^ cells per ml) were stimulated with rhIL-1β (10ng/ml) for 3 hours, with or without the pretreatment with IL-1Ra (10 μg/ml). Cell extracts were immunoblotted for IκBζ with anti-IκBζ antiserum (1:2000). IκBζ expression was normalized to protein loading control and represented as densitometric units. Error bars represent the mean ± SEM of 3 independent experiments. NT stands for no treatment control. Beta actin was used as protein loading control for both (A) and (B).

## DISCUSSION

Various groups have explored the role of IκBζ in regulating respiratory immunity. IκBζ has been demonstrated to modulate the expression of pro-inflammatory cytokines like GMCSF and IL-6 and anti-microbial peptides like NGAL in airway epithelial cells that line the respiratory mucosal surface (4, 5, 7, 16). This is relevant to lung defense because GMCSF is critical for the maintenance of lung surfactant homeostasis (18) and protection from pneumonia (19). *Nfkbiz* knockout mice are severely deficient in GMCSF (15). Recently, we demonstrated IκBζ’s role in regulating inflammatory responses to pneumococcus infection (20), further emphasizing the importance of IκBζ in lung immune defense. Original studies reported that mononuclear phagocytes which exist in the pristine vascular environment require additional TLR or IL-1R agonists to induce IκBζ expression (8). However, constitutive expression of IκBζ messenger RNA levels in epithelial cells lining various mucosal surfaces, including the eyes and the skin have been reported (13, 14). These findings suggest the likelihood that local recognition of PAMPs in exposed tissues can induce a steady state expression of IκBζ. However, here for the first time, we provide strong evidence for constitutive expression of IκBζ protein in airway epithelium, in the absence of exogenous agonist activity. This unique expression of IκBζ has been consistently observed in all of our previous studies in which we used immunoblotting to study IκBζ expression in primary respiratory epithelial cells cultured in the absence of known PAMPs and IL-1 (7, 20).

Why recombinant IκBζ-S blocked all the nonspecific bands, in addition to the IκBζ-L band in **figure 3B** is unclear. However, it is noteworthy that it is the same purified rIκBζ-S which was the antigen against which the anti-IκBζ antiserum was generated in our laboratory. It is hence plausible that the non-specific protein fragments that may have sequence similarities with the recombinant protein also get detected by the polyclonal antibody population in the antiserum and as a result, get blocked by the recombinant protein.

The other protein detected by the anti-IκBζ antiserum, lamin B1, predominantly localizes in the nucleus, where functional IκBζ also accumulates. This made the interpretation of our immunostaining data difficult. Testing our anti-IκBζ antiserum on airway tissue from mice with conditional complete *Nfkbiz* knockout in the lungs will be needed to absolutely confirm specificity. Our data highlights the need for a highly sensitive and specific IκBζ antibody, especially for use in IHC-based detection of the protein in tissue samples. This study also emphasizes the importance of having the right controls to prove antibody specificity while demonstrating IκBζ expression in tissues using IHC.

Our observation concurs with the known constitutive expression of IκBζ in other mucosal surfaces such as skin and eyes, further supporting our findings. IκBζ’s constitutive expression in airway epithelium suggests a function and regulation that may differ from its classical TLR/IL-1 induction seen in immune cells. Interestingly, the baseline IκBζ expression had no apparent function in terms of activating cytokine release from primary human bronchial epithelial cells (7). It is conceivable that IκBζ, like other members of the IκB family, requires post-translational modification such as phosphorylation, as an essential step for its functional activation. Smearing of the IκBζ-L band (**figure 5A**) and its pattern with a slight increase in size of isoforms in the acidic range of the 2D gel (**figure 4A**) are suggestive of the same. Nevertheless, IL-1 was able to induce further IκBζ protein beyond the baseline expression which was associated with subsequent cytokine release in these cells (7). Future studies evaluating IκBζ’s signal specific interactions with its known partners like NFκB or akirin2 (21) or its post-translational modification as an activation step, may be able to identify the exact function of the uniquely expressed IκBζ in lung epithelium. Nevertheless, our current data support a vital role for IκBζ in maintaining homeostasis of lung immunity.

## CONCLUSION

In conclusion, our data demonstrate constitutive IκBζ protein expression in the airway epithelium of both humans and mice. This constitutive expression likely is critical to lung host defense via effects on downstream transcriptional targets. Understanding how this constitutive expression of IκBζ changes during disease conditions will help evaluate the protein as a potential biomarker and a therapy candidate. This study also highlights the need for a highly sensitive and specific antibody for IκBζ detection, especially for use in immunohistochemistry.

## DECLARATIONS

### Ethics approval and consent to participate

All animal experiments were performed according to animal protocols approved by the Animal Care Use Committee of the Ohio State University College of Medicine. Anonymized human lung tissue slides were provided by the Ohio State University Department of Pathology, Tissue Archive Service. Other investigators may have received specimens from the same subjects.

### Availability of data and materials

All data generated or analyzed during this study are included in this published article [and its supplementary information files]

### Competing interests

The authors declare that they have no competing interests.

### Funding

This work was supported by NIH grants HL089440 and HL076278 to Dr. Wewers.

### Author Contributions

KS^1^ and MDW conceived and designed experiments;

KS^1^ and MDW analyzed and interpreted data;

KS^1^ and MDW drafted the manuscript for important intellectual content;

KS^1^ and SM performed experiments;

KSO and SL provided lung tissue from *Nfkbiz^−/−^* knockout mice;

PNB and HS provided lung tissue from gnotobiotic mice;

AS provided lung tissue from normal wild type mice;

KS^6^ provided human lung tissue slides.

All authors reviewed the final manuscript.

## Acknowledgements

We thank Masami Morimatsu (Sapporo, Japan) for kindly providing us the *Nfkbiz^−/−^* knockout mice. We thank Arun Tiwari, a former member of OSU’s proteomics facility, for his help with generating the mass spectrometry data. We thank Florinda Jaynes and Alan Flechtner from the Comparative Pathology and Mouse Phenotyping Shared Resource at OSU, for performing IHC on our samples. We thank Sudarshan Seshadri and Yashaswini Kannan for establishing the tools to study IκBζ in our laboratory.

---

**Supplementary Figure S1. Anti-IκBζ antiserum detects IκBζ protein. (A)** BEAS2B cells were (1.5×10^5^ cells/ml) transfected with 100pmol of scrambled siRNA control or siRNA specific to IκBζ and stimulated with rhIL-1β (10ng/ml) for 3 hours. Cell lysates were immunoblotted for IκBζ with anti-IκBζ antiserum (1:2000). **(B)** Extracts from BEAS2B cells stimulated with IL-1β were immunoblotted with IκBζ antibodies from Thermo Fisher Scientific (PA5-52703), Sigma Aldrich (HPA010547), Abcam (ab221914) and LS Biosciences (LS-C294627) at their recommended dilutions or our laboratory-made anti-IκBζ antiserum (1:2000). Beta actin was used as equal protein loading control for both (A) and (B). Asterisk represents non-specific proteins detected. The results are representative of at least 3 independent experiments.

**Supplementary Figure S2. IκBζ expression in trachea from gnotobiotic mice**. Tracheal slices from gnotobiotic mice were immunostained with secondary antibody alone (2° ab) (1:1000), IgG isotype control (1:1500), or anti-IκBζ antiserum (1:1500), observed at 100, 400 and 1000X magnifications. The scale bar denotes 50μm for the 1000X magnification panels. The results are representative of 3 different mice. Red boxes represent regions that were captured at higher magnifications.

**Supplementary Figure S3. THP-1 nuclear extract blocks non-specific IgGs in anti-IκBζ antiserum. (A)** Immunoblot of non-specific lamin B1 protein from the nuclear extract (NE), cytoplasmic extract (CE) and nuclear pellet (Nu) of unstimulated THP-1 cells, using Lamin B1 antibody at its recommended dilution. IL-1β stimulated BEAS2B cell extracts **(B)** immunoblotted with our laboratory made anti-IκBζ antiserum alone (1:5000) or with the anti-IκBζ antiserum pre-cleared with excess unstimulated THP-1 NE and (C) immunostained with secondary antibody only (2° ab) (Panel 1), IgG isotype control (Panel 2), anti-IκBζ antiserum (1:5000) (Panel 3) or anti-IκBζ antiserum pre-cleared with excess THP-1 NE (Panel 4) and imaged at 100X magnification. NT stands for no treatment. The results are representative of 7-8 independent experiments.

