## Supplementary figures and images for "IκBζ is constitutively expressed in human and murine airway epithelium"

### supl fig 1

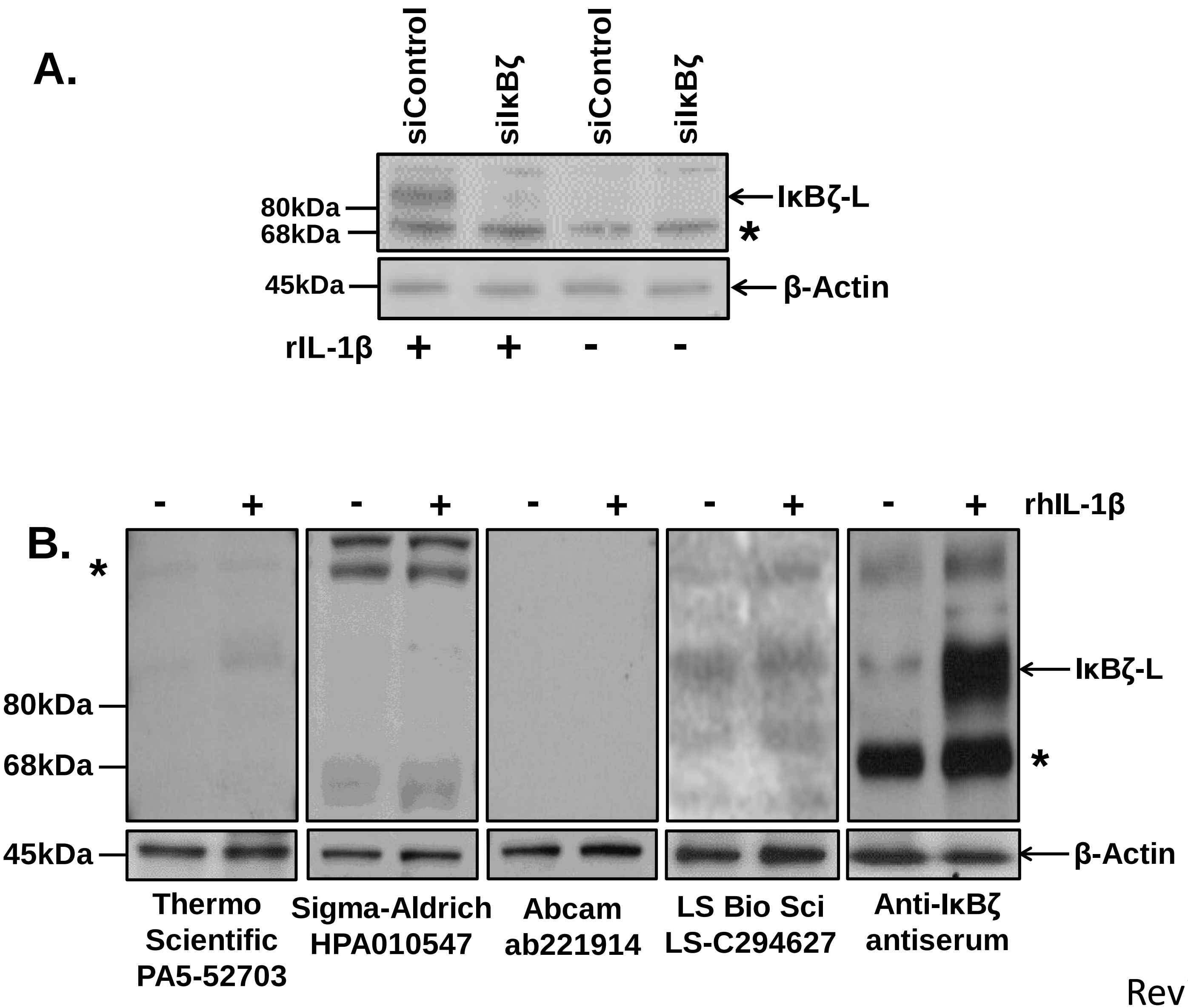


Revised
Sup Fig 1

### supl fig 2

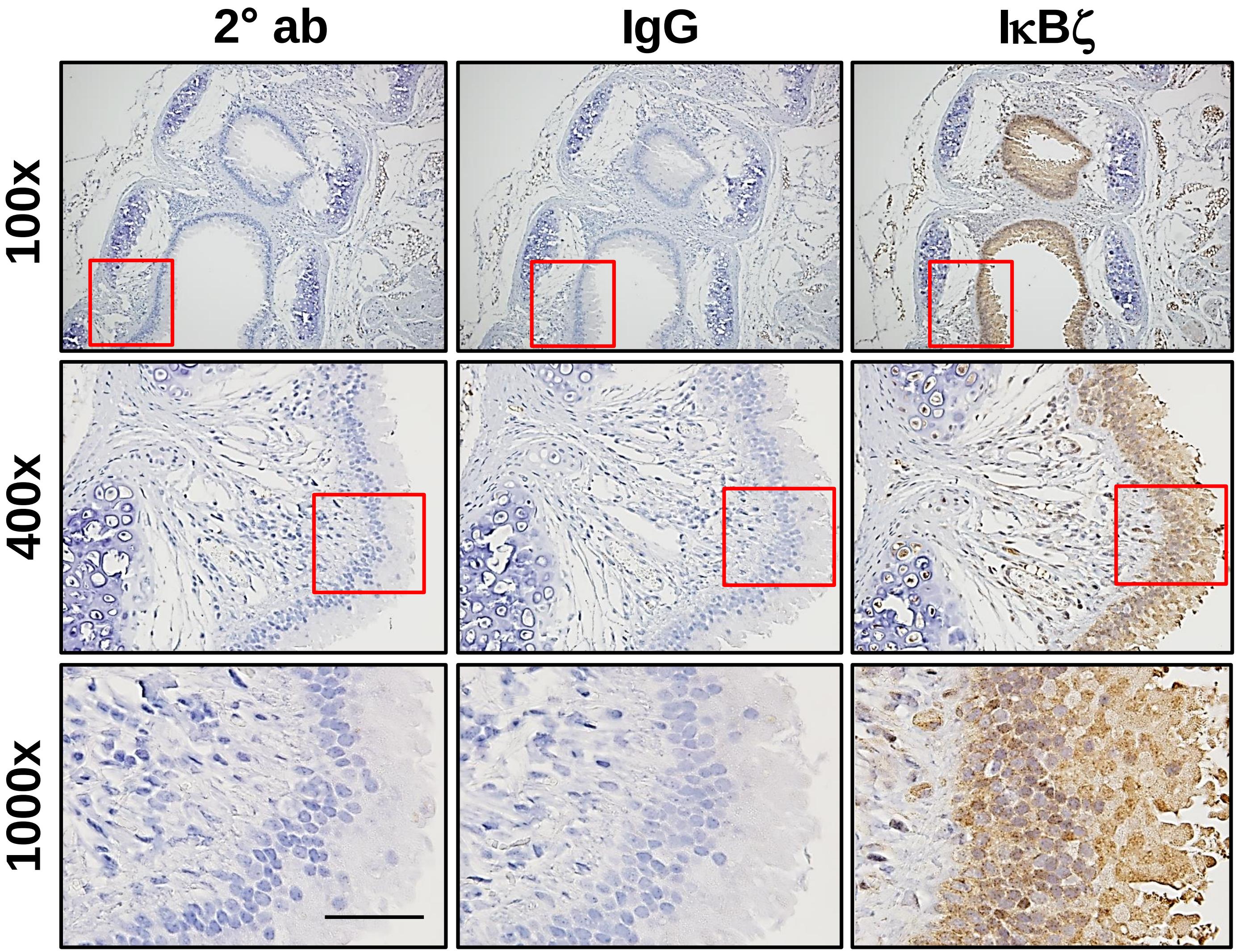


Revised
Sup Fig 2

### supl fig 3

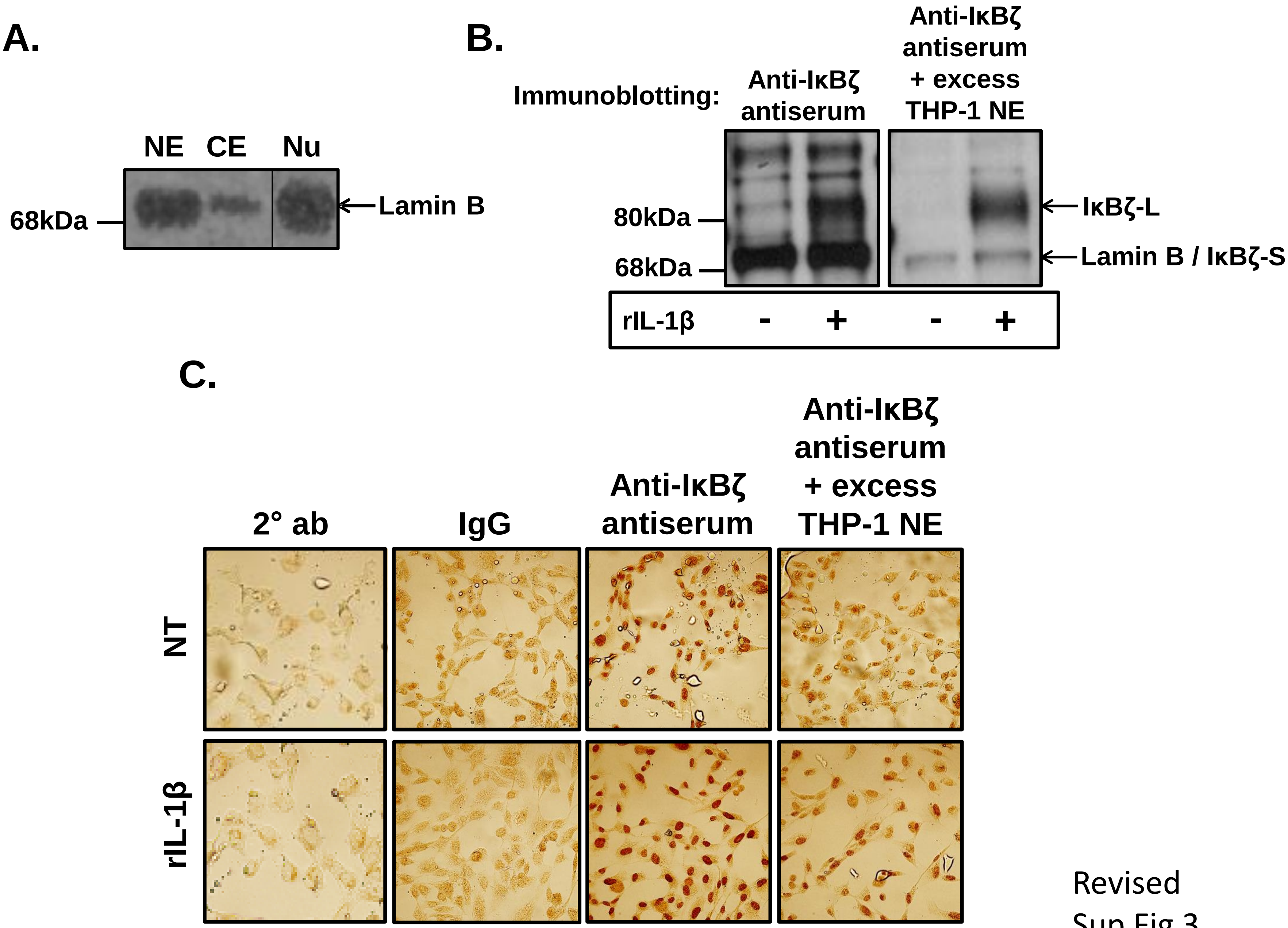


Revised
Sup Fig 3
